# Identification of repurposable cytoprotective drugs for Vanishing White Matter Disease

**DOI:** 10.1101/2020.06.02.131052

**Authors:** Neville Ng, Mauricio Castro Cabral-da-Silva, Simon Maksour, Tracey Berg, Martin Engel, Dina M. Silva, Dzung Do-Ha, Jeremy S. Lum, Sonia Sanz Muñoz, Nadia Suarez-Bosche, Claire H. Stevens, Lezanne Ooi

## Abstract

Vanishing white matter disease (VWMD) is a rare leukodystrophy involving loss of function mutations of the guanine exchange factor eIF2B and typically presenting with juvenile onset. We aimed to identify repurposable FDA approved drugs in an *in vitro* drug screen using patient-derived fibroblasts and induced pluripotent stem cell (iPSC)-derived astrocytes. Dysregulated GADD34 and CHOP were identified in patient fibroblasts and iPSC-derived astrocytes under proteasomal stress conditions. A drug screen from a 2400 FDA approved drug library with *EIF2B5* disease patient fibroblasts identified 113 anti-inflammatory drugs as a major class of hits with cytoprotective effects. A panel of potential candidate drugs including berberine, deflazacort, ursodiol, zileuton, guanabenz and Anavex 2-73, and preclinical ISRIB, increased cell survival of MG132-stressed *EIF2B2* and *EIF2B5* disease VWMD astrocytes, and were further investigated for their effect on the integrated stress response and mitochondrial stress. ISRIB but not other drugs significantly affected eIF2α phosphorylation and GADD34 expression. Ursodiol demonstrated capacity to reduce complex I subunit upregulation, ameliorate oxidative stress, loss of mitochondrial membrane potential and upregulation of eIF2B subunits in VWMD astrocytes, highlighting its potential as a cytoprotective compound for VWMD.

## Introduction

Vanishing white matter disease (VWMD) is a rare, autosomal recessive leukodystrophy, caused by mutations in the genes, *EIF2B1, EIF2B2, EIF2B3, EIF2B4, EIF2B5,* encoding the eukaryotic initiation factor eIF2B (1). The eIF2B protein is a guanine nucleotide exchange factor that is involved in the integrated stress response (ISR) and loss of function mutations in both alleles of an *EIF2B* gene leads to VWMD. VWMD is a debilitating and progressive disease; patients are often diagnosed as children and survive few years beyond diagnosis. Known mutations of *EIF2B* are considered to cause loss of function of the wild-type protein, with variable onset, progression and severity, dependent on the precise mutations and environmental stress factors (2).

The eIF2B proteins regulate mRNA translation, converting the inactive eIF2-GDP to the active eIF2-GTP form (3). Activation the cytoprotective ISR leads to phosphorylation of eIF2α (p-eIF2α), binding to eIF2B, translational repression, and upregulation of stress-induced genes (4). These include GADD34, which facilitates dephosphorylation of p-eIF2α toward recovery from stress and resumption of normal protein translation in a negative feedback loop, and transcription factors ATF4 and CHOP (5). The partial loss of function of eIF2B can lead to delayed translation of stress-induced genes and dysregulated ISR expression (2). Although eIF2B is ubiquitously expressed and plays a role in multiple cell types, the disease manifests most significantly in the loss of white matter of the brain (6).

*EIF2B* mutant mouse models and induced pluripotent stem cell (iPSC) models (7, 8) have identified a central role for dysfunctional astrocytes in the development of VWMD with evidence for astrocytic apoptosis (8) and an inability to promote oligodendrocyte maturation (6). A key driver of cellular pathogenesis in VWMD involves the ISR, with alterations in responses to endoplasmic reticulum stress, proteasomal stress and oxidative stress (3). Mitochondrial dysfunction and upregulated reactive oxygen species have been identified to be upregulated in *EIF2B5*^R132H/R132H^ murine fibroblasts and astrocytes (9). Currently there are no approved treatments for VWMD, hence the aim of this research was to identify candidate drugs from an FDA approved drug library that could protect VWMD patient cells against cellular stressors relevant to the disease.

## Results and Discussion

### VWMD patient fibroblasts and astrocytes exhibit dysregulated ISR marker expression

The partial loss of function of eIF2B has been observed to suppress both global and stress-induced protein translation, in response to ER stress in VWMD patient lymphoblasts (2, 10). Given that eIF2B is a ubiquitously expressed protein we anticipated that a dysfunctional ISR may be evident in patient fibroblasts and iPSC-derived astrocytes. Fibroblasts were reprogrammed into iPSCs from two VWMD patients, along with gender-matched relatives as non-disease controls (Figure S1). The VWMD1 iPSC line was generated from a patient bearing mutations in the *EIF2B5* gene (encoding eIF2Bε^R133H/A403V^), whilst the VWMD6 iPSC line was generated from a patient bearing mutations in the *EIF2B2* gene (encoding eIF2Bβ^G200V/E213G^). A previous study identified that white matter-derived astrocytes, generated using a CNTF-based differentiation protocol, showed a more vulnerable phenotype to stress, compared to grey matter astrocytes generated using FBS (7). Consequently, we generated astrocytes from iPSCs using a CNTF-based method (Figure S2).

Responses to oxidative, proteasomal and endoplasmic reticulum (ER) stresses are all proposed to be affected in VWMD cell and animal-based models (2, 11). Thus, we evaluated the dose-dependent effect of H_2_O_2_, MG132 and thapsigargin stressors, as mediators of oxidative, proteasomal and ER stress, respectively (Figure S3). There was a significant reduction in cell survival for VWMD fibroblasts and iPSC-derived astrocytes, compared to non-disease controls, under all three stressors (Figure S3).

MG132 is commonly utilised as a proteasomal inhibitor that can also induce ER and oxidative stress via the unfolded protein response, all of which trigger the ISR (12). A key consequence of partial loss of eIF2B function is disruption of ISR homeostasis, which can be assessed by measuring the expression levels of ISR-relevant markers. The control of eIF2α phosphorylation or dephosphorylation acts as a pivotal mechanism that regulates global protein synthesis in response to cell stress, as part of the ISR (13); GADD34 dephosphorylates eIF2α (14). The transcription factor ATF4 is an activator of the ISR in response to stress, while CHOP is activated by the ISR to promote apoptosis (15). The expression levels of these four ISR-relevant proteins, p-eIF2α (normalised to eIF2α), GADD34, ATF4 and CHOP, were compared in VWMD and control fibroblasts and iPSC-derived astrocytes under MG132 stress (Figure 1). Under proteasomal stress, VWMD and control fibroblasts exhibited similar decreases in eIF2α phosphorylation and GADD34 expression (Figure 1A-D), consistent with VWMD lymphoblasts (2). However, elevated levels of ATF4 and CHOP were observed in MG132-stressed VWMD fibroblasts, compared to controls, suggesting increased activation of the ISR. MG132-stressed VWMD fibroblasts and astrocytes also exhibited increased CHOP, while astrocytes exhibited increased GADD34 (Figure 1E-H), consistent with glia in animal model studies (10).

**Figure 1.**
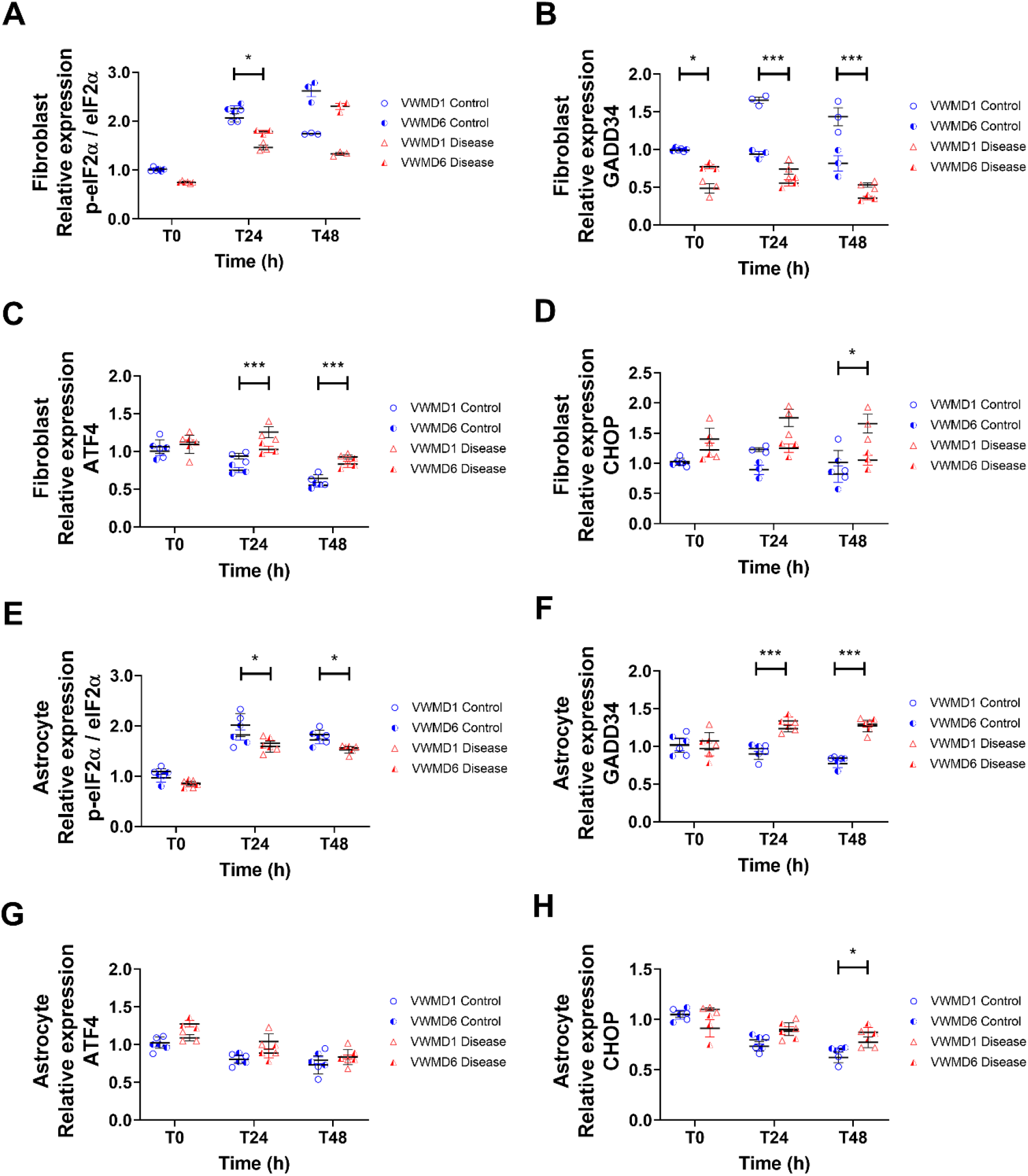
ISR marker protein expression is affected in VWMD disease fibroblasts and astrocytes. (A-H) Effect of MG132 stress on protein expression of the ISR markers, p-eIF2α (normalised to eIF2α), GADD34, ATF4 or CHOP, at 0, 24 or 48 h incubation period. Protein levels were quantified by western blot, normalised to total protein, and are shown relative to control at time 0. Individual data points are shown from VWMD1 patient *EIF2B5*^R133H/A403V^ (open red triangle); VWMD6 patient *EIF2B2*^*G200V/E213G*^ (closed red triangle) or their non-disease controls VWMD1 Control *EIF2B5*^wt/wt^; VWMD6 Control *EIF2B2*^*G200V/wt*^ with mean ± SEM, n = 3. Significant differences were identified by two-way ANOVA followed by Holm-Sidak posthoc test, * p < 0.05, ** p < 0.01, *** p < 0.001. Representative blots shown in (Figure S3).

### Cytoprotective drug screen in VWMD patient lines

Based on the established capacity of the proteasomal inhibitor, MG132, to induce proteasomal stress, oxidative stress, and exacerbate ISR disease phenotypes, we performed a first-pass drug screen for candidates able to protect against the effect of MG132 in VWMD1 *EIF2B5*^R133H/A403V^ patient fibroblasts, with cell viability assessed by resazurin reduction activity (Figure 2).

**Figure 2.**
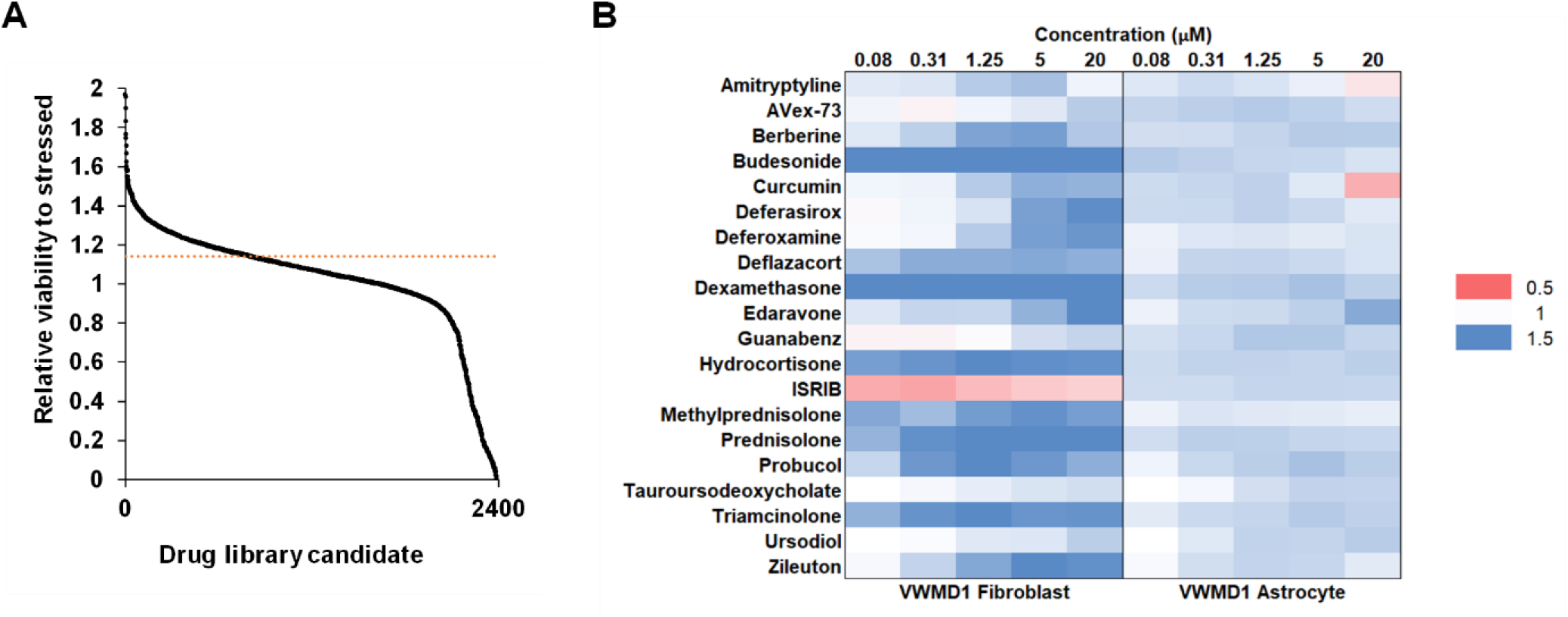
Drug screen of cytoprotective candidates. (A) VWMD1 *EIF2B5*^R133H/A403V^ disease fibroblasts were coincubated with MG132 and each of 2400 drugs from an FDA-approved drug library. Cell viability was measured and normalised to cell viability with MG132 stressor in the absence of drug. Drugs were ranked based on increase in cell viability (n = 3). Cytoprotection ranged from 2-fold increase in viability to 0 (100% cell death for cytotoxic drugs). Horizontal dotted red line indicates 1.5 x standard deviation of MG132 stressed controls. (B) Hit candidates were taken forward to assess their dose response effect on cell viability of VWMD1 *EIF2B5*^R133H/A403V^ fibroblasts and iPSC-derived astrocytes, relative to MG132 stress (n = 5-6). ISRIB was included due to its previous identification as a potential treatment for VWMD, although it caused significant toxicity in fibroblasts. Heat map represents cytoprotection (blue) versus cytotoxicity (red) of individual drugs.

Following the initial drug screen, a panel of 20 compounds (Table S2) was selected for downstream evaluation in VWMD1 *EIF2B5*^R133H/A403V^ fibroblasts and in both VWMD1 *EIF2B5*^R133H/A403V^ and VWMD6 *EIF2B2*^*G200V/E213G*^ patient-derived astrocyte lines. The compounds selected for downstream assays were chosen based on their translational premise, including considerations of bioavailability, route of administration and complications in treatment. The drug panel included 15 protective compounds from the screen (>1.5 × standard deviation of MG132-stressed controls), and a further five compounds with relevant modes of action for VWMD. Overall, this panel included glucocorticosteroids, bile acids, iron chelators, antioxidants, ISR modulators and sigma-1 receptor agonists. Nominated drugs included guanabenz, ISRIB, sigma-1 receptor agonists, AVex-73 and amitriptyline, tauroursodeocycholic acid and alkaloid berberine. All candidates elicited cytoprotective effects against MG132-induced stress at varying concentrations in VWMD1 fibroblasts, with the exception of ISRIB. Ursodiol and its taurine derivative, tauroursodeoxycholic acid, showed similar cytoprotective efficacies. We observed that anti-inflammatories were amongst the largest category of compounds that improved cell viability under proteasomal stress, with a high proportion of these being glucocorticosteroids. Steroids can regulate inflammation, mitochondrial function and apoptosis toward neuroprotective effect in brain injury, Alzheimer’s disease, Parkinson’s disease, multiple sclerosis, and stroke (16). Although glucocorticosteroids are a commonly administered class of drugs, an anecdotal study of corticosteroids on three VWMD patients did not identify benefits and the patients were removed from this treatment due to potential clinical complications (17). Deflazacort was selected as a representative glucocorticosteroid on the basis of fewer reports of adverse effects in the literature and its use in the clinic, including in children with muscular dystrophy (18, 19). The recent demonstration of mitochondrial dysfunction and inefficient respiration in murine models of VWMD (9) has expanded the search for possible therapeutics to include mitochondrial protective compounds and antioxidants (3, 9). The protective effect of the antioxidant edaravone was evident in astrocytes at a higher efficacy than any other candidate, potentially due to its well established radical scavenging activity (20). Edaravone was recently approved for amyotrophic lateral sclerosis and while its administration is currently limited to intravenous injection, oral and mucosal formulations are in development (21–23). The sigma-1 receptor is a chaperone protein in ER membranes that governs a range of cellular processes, including calcium homeostasis and reactive oxygen species accumulation (11). However, in this study, the cytoprotective effect of the sigma-1 receptor agonists, AVex-73 and amitriptyline, was limited at higher concentrations (≥5 μM).

VWMD6 *EIF2B2*^G200V/E213G^ iPSC-derived astrocytes confirmed the findings in VWMD1 *EIF2B5*^R133H/A403V^ iPSC-derived astrocytes (Figure S4). Of those tested, the majority of the drugs that were protective in VWMD1 fibroblasts were also protective in VWMD1 and VMWD6 astrocytes, with the exception of amitryptiline, curcumin and budesonide (Figures 2, S4).

### Effect of candidate drugs on mode of cell death

We further investigated the effects of a panel of compounds, in order to provide insight into potential modes of cytoprotective action. For these experiments we assessed drugs with strong translational potential for VWMD (Figures 2, S4), including Avex-73, berberine, deflazacort, guanabenz, ISRIB, ursodiol and zileuton. We did not include edaravone, despite its promising effects in patient fibroblasts and astrocytes, due to its intravenous route of administration.

Assays to investigate potential drug mechanisms were performed using MG132 stress-induced VWMD1 *EIF2B5*^R133H/A403V^ patient astrocytes, as the most common VWMD subunit mutation (24). To determine whether candidate drugs reduced the proportion of bulk cell death, the membrane permeabilisation of treated astrocyte cultures was evaluated. In MG132-stressed VWMD1 astrocytes there were no significant changes in the percentage of membrane-permeablised cells, with the exception of ISRIB increasing the proportion of membrane-permeablised cells (Figure 3A). To test whether the drugs caused a reduction in mitochondrial apoptosis, the *BAX*:*BCL2* ratio was evaluated. There was no detectable change in *BAX:BCL2* ratio caused by any of the drug candidates, with the exception of an increase in *BAX:BCL2* caused by ISRIB (Figure 3B). Together these data show that astrocytes coincubated with MG132 and ISRIB exhibited an overall increase in cell viability, despite having an elevated proportion of membrane-permeabilised cells and increased *BAX:BCL2* ratio. This apparent paradox may reflect the ability of ISRIB to promote protein synthesis under stress conditions, and thus support a greater turnover of the cell population. ISRIB may thus have an adverse effect on cell survival under acute stress conditions, particularly for non-cycling cells, such as neurons. The ramifications of this are yet to be reported in animal studies (25, 26), however the enhanced effect of cytotoxic drugs under an ISRIB-suppressed ISR have been recently demonstrated in prostate and pancreatic cancer studies (27, 28). Further testing of ISRIB in animal studies will uncover details of its precise role on cell survival during different stress conditions.

**Figure 3.**
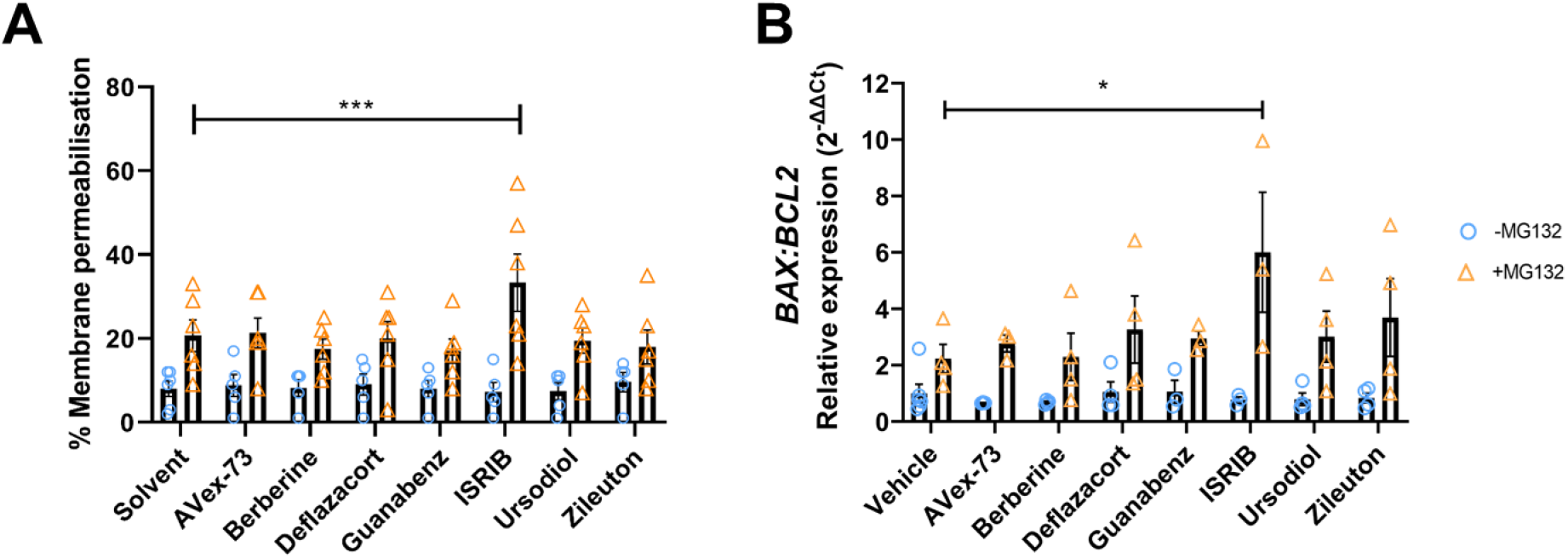
Effect of candidate drugs on membrane permeabilisation and mitochondrial pathway cell death markers in vehicle control (blue circle) or 0.1 μM MG132-treated (orange triangle) VWMD1 *EIF2B5*^R133H/A403V^ patient iPSC-derived astrocytes. Cells were treated with candidate drugs, with or without MG132 for 24 h. (A) Ethidium homodimer was used to quantify % membrane permeabilised astrocytes (n = 5-6). (B) RT-qPCR was used to measure gene expression changes of the mitochondrial apoptosis pathway indicator, *BAX*:*BCL2* ratio (n = 3-4). Individual data points are shown with mean ± SEM; significant differences are shown by * p < 0.05, *** p < 0.001, identified by two-way ANOVA followed by Holm-Sidak posthoc test.

### Effect of candidate drugs on ISR-relevant proteins, p-eIF2α, GADD34, ATF4 and CHOP

A key consequence of partial loss of eIF2B function is disruption of ISR homeostasis, thus candidate drugs were assessed for their effect on the ISR under MG132 stress. The expression levels of the ISR-relevant proteins, p-eIF2α (normalised to eIF2α), GADD34, ATF4 and CHOP, were evaluated following candidate drug treatment (Figure 4A-D). ISRIB markedly increased p-eIF2α and decreased GADD34 and CHOP expression under MG132 stress conditions (Figure 4A-D). No significant changes in ATF4 were detected by any of the drugs (Figure 4C). However, AVex-73, berberine and deflazacort significantly decreased CHOP expression in the presence of MG132 (Figure 4D).

**Figure 4.**
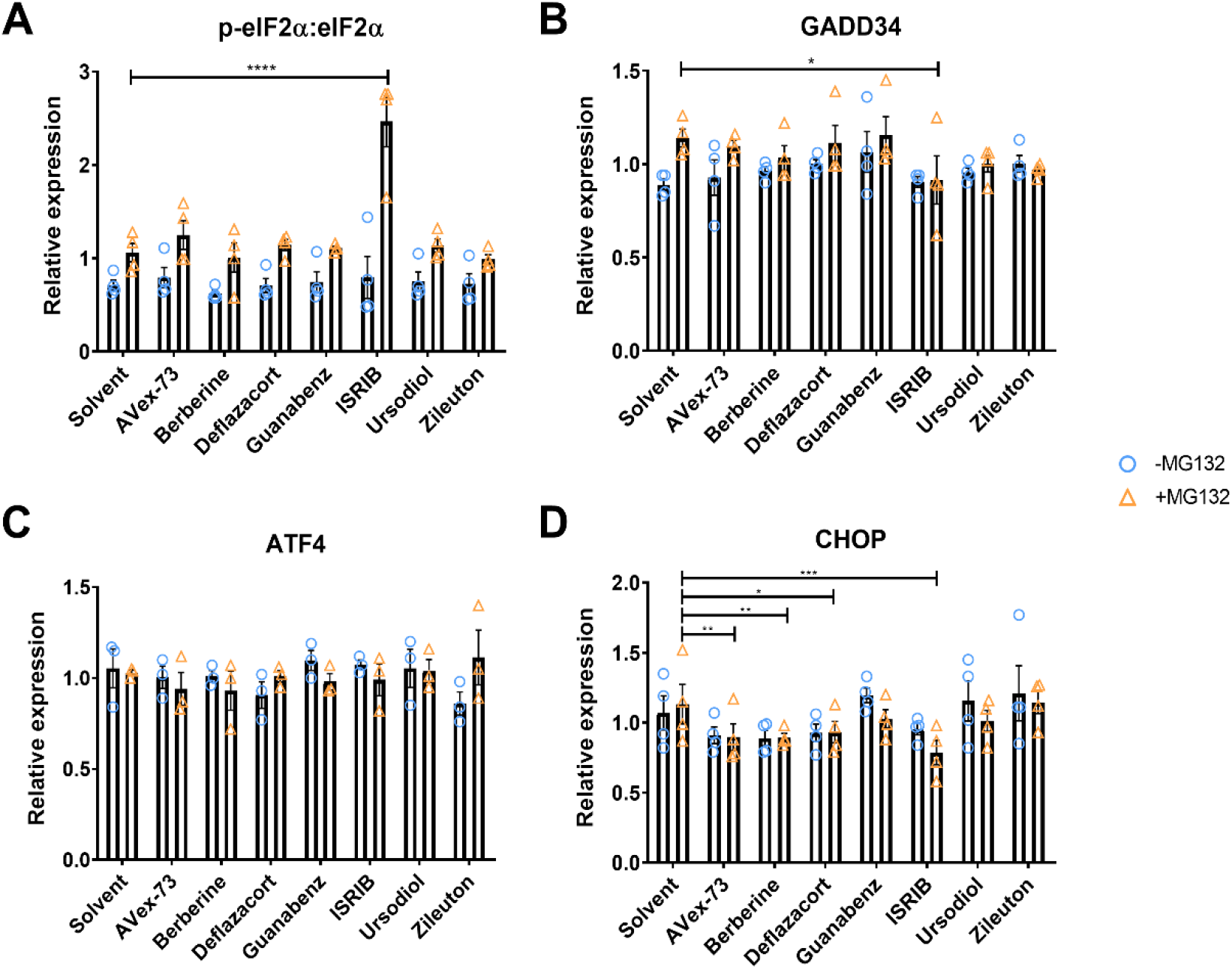
Effect of candidate drugs on ISR markers in vehicle control (blue circle) or 0.1 μM MG132-treated (orange triangle) VWMD1 *EIF2B5*^R133H/A403V^ patient iPSC-derived astrocytes. Cells were treated with candidate drugs, with or without MG132 for 24 h. Protein expression was measured by western blot and normalised to total protein for the ISR markers: (A) phosphorylated eIF2α, normalised to eIF2α; (B) GADD34; (C) ATF4; (D) CHOP. Individual data points are shown with mean ± SEM (n=4). Significant differences were identified by two-way ANOVA, followed by Holm-Sidak posthoc test * p < 0.05, ** p < 0.01, *** p < 0.001, **** p < 0.0001. Representative western blots shown in (Figure S6).

Previous cell stress and neurodegeneration studies have established ISR-modulating roles for guanabenz and ISRIB (29, 30), hence their inclusion in our panel of compounds. Guanabenz is considered to exert cytoprotective effects by inhibiting the activity of GADD34 to recruit eIF2 phosphatases, thus prolonging translation inhibition and avoiding the stress of resuming protein synthesis (29). However, at concentrations of guanabenz that induced a cytoprotective effect (5 μM) we did not observe a significant impact on expression of any ISR markers. Conversely, ISRIB-mediated cytoprotection of astrocytes (1.25 μM) was associated with increased eIF2α phosphorylation, and downregulation of GADD34 and CHOP.

ISRIB is thought to allow eIF2B to escape inhibitory complex formation with p-eIF2α, in order to limit repression of protein synthesis while under ER stress (10, 31). An analogue of ISRIB was recently shown to improve ISR signature, motor function and myelin loss in a *EIF2B5* VWMD mouse model (10, 31). However, in this study, ISRIB exacerbated MG132 toxicity in *EIF2B5* VWMD fibroblasts at all doses (0.08-20 μM), consistent with prior reports that ISRIB enhances thapsigargin-induced apoptosis in HEK293T cells (26). It is also worth noting that the stabilising effect of ISRIB may be mutation-dependent (3). The phytochemical, berberine, was included in the panel because it is considered to be a well-tolerated natural compound (32). In MG132-stressed astrocytes berberine treatment increased cell viability, potentially via decreasing pro-apoptotic CHOP (Figure 4D). The antioxidant, anti-inflammatory, ER and mitochondrial protective effects of berberine have been noted in numerous studies, including increased levels of antioxidants, superoxide dismutase and glutathione, inhibition of caspase 3 activity and apoptosis and decreased cytochrome C and *BAX:BCL2* ratio in ischemic injury and diabetic animal models (33, 34).

### Effect of candidate drugs on indicators of mitochondrial function

The eIF2B mutations in murine models have been shown to affect mitochondrial complex I function and increase mitochondrial abundance (35). To identify potential mitochondrial protective mechanisms, we assessed the effect of the candidate drugs on the expression levels of mitochondrial complex I using the markers, *NDUFA8* and *NDUFS7,* in VWMD1 *EIF2B5*^R133H/A403V^ astrocytes. Ursodiol significantly prevented complex I subunit increases in *NDUFS7* caused by MG132, while AVex-73 and deflazacort reduced *NDUFA8* expression (Figure 5A-B). A reduction in complex I subunit expression and CHOP expression may reflect the ability of deflazacort to improve mitochondrial biogenesis (36). AVex-73 decreased VWMD1 astrocyte expression of complex I subunit in the absence of stressor, which may indicate a decrease in respiratory burden, although this was not evident under MG132 stress (11). Mutations in eIF2B genes impair mitochondrial function during oxidative stress conditions and suggest a dysregulated ISR in VWMD murine fibroblasts and astrocytes (9), consistent with our findings.

**Figure 5.**
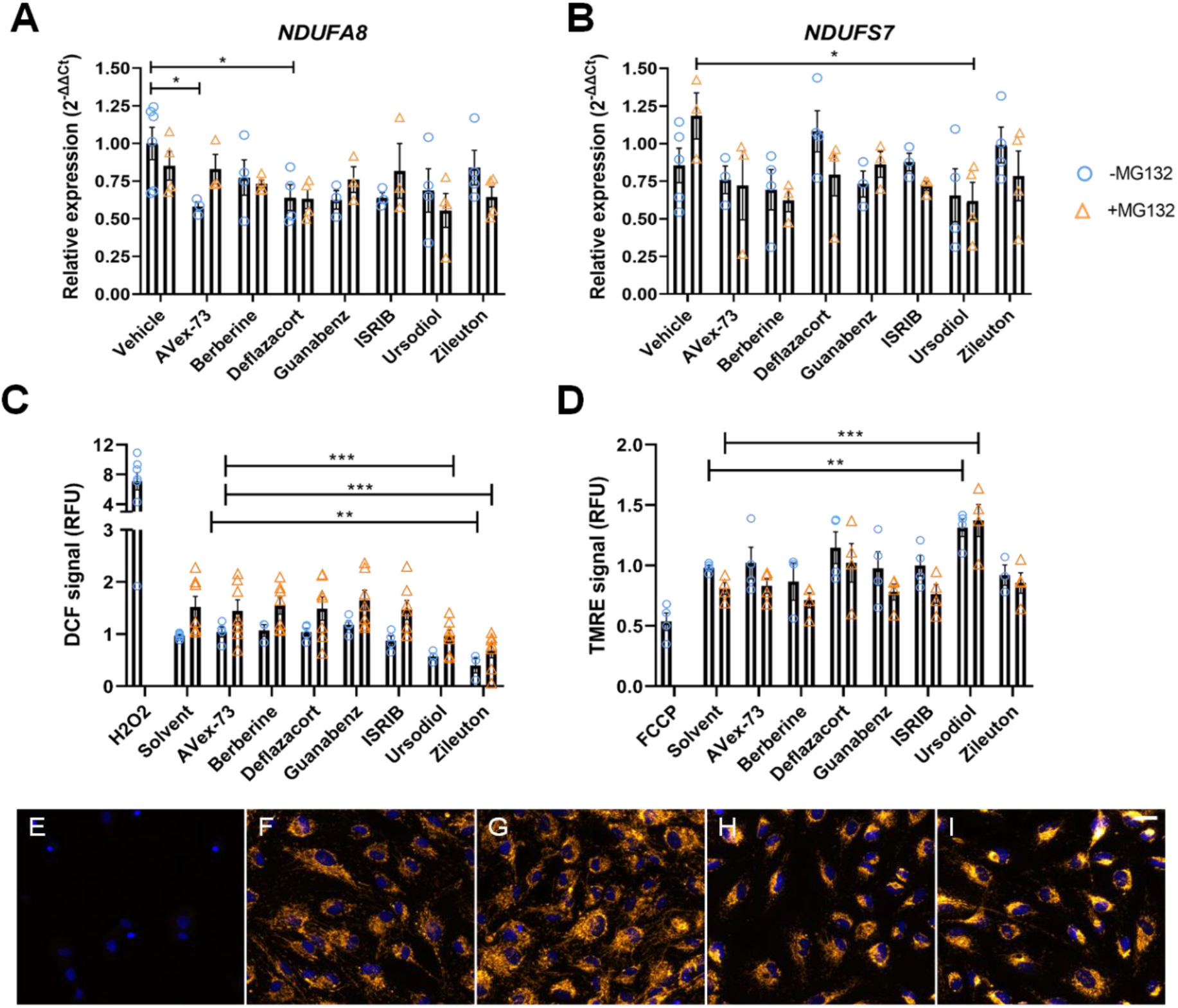
Effect of candidate drugs on complex I subunit expression, oxidative stress and mitochondrial membrane potential in VWMD1 *EIF2B5*^R133H/A403V^ disease astrocytes. VWMD1 iPSC-derived astrocytes were treated with vehicle control (DMSO; blue circle) or 0.1 μM MG132 (orange triangle) and candidate drugs for 24 h. Expression of mitochondrial complex I subunits was measured by RT-qPCR for (A) *NDUFA8* and (B) *NDUFS7*. (C) Reactive oxygen species was measured via DCF fluorescence and (D) TMRE fluorescence was used to measure mitochondrial membrane potential. Representative confocal fluorescence microscopy images of: (E) FCCP (protonophore, mitochondrial uncoupler, loss of TMRE fluorescence), (F) TMRE assay solvent, (G) ursodiol, (H) MG132, (I) MG132 + ursodiol (scale bar = 30 μm) (orange = TMRE; blue = Hoechst 33342 nuclear stain). For A-D individual data points are shown and mean ± SEM from replicate differentiations (n=4-6); significant differences were identified by two-way ANOVA followed by Holm-Sidak posthoc test * p < 0.05, ** p < 0.01, *** p < 0.001.

The candidate drugs were further investigated for their effects on oxidative stress and mitochondrial membrane potential. Ursodiol and zileuton significantly decreased DCF signal, as an indicator of reactive oxygen species generation, in VWMD1 *EIF2B5*^R133H/A403V^ patient astrocytes, in the presence or absence of MG132 stress (Figure 5C). Of the anti-inflammatory candidate drugs, the 5-lipoxygenase antagonist zileuton, demonstrated significant reduction in generation of reactive oxygen species, a finding that correlates with its radical scavenging activity (37). Ursodiol also increased relative mitochondrial membrane potential of VWMD1 *EIF2B5*^R133H/A403V^ astrocytes under both unstressed and stressed conditions, based on the TMRE assay (Figure 5D-I). Together, these data suggest that ursodiol may promote mitochondrial function and reduce oxidative stress in VWMD astrocytes.

### Ursodiol rescues repression of *EIF2B2* and *EIF2B5* gene expression

The potential cytoprotective effect of candidate drugs, as a result of upregulation of *EIF2B* genes, was considered. In MG132-stressed VWMD1 *EIF2B5*^R133H/A403V^ astrocytes, ursodiol rescued reductions in *EIF2B2* and *EIF2B5* expression (Figure 6).

**Figure 6.**
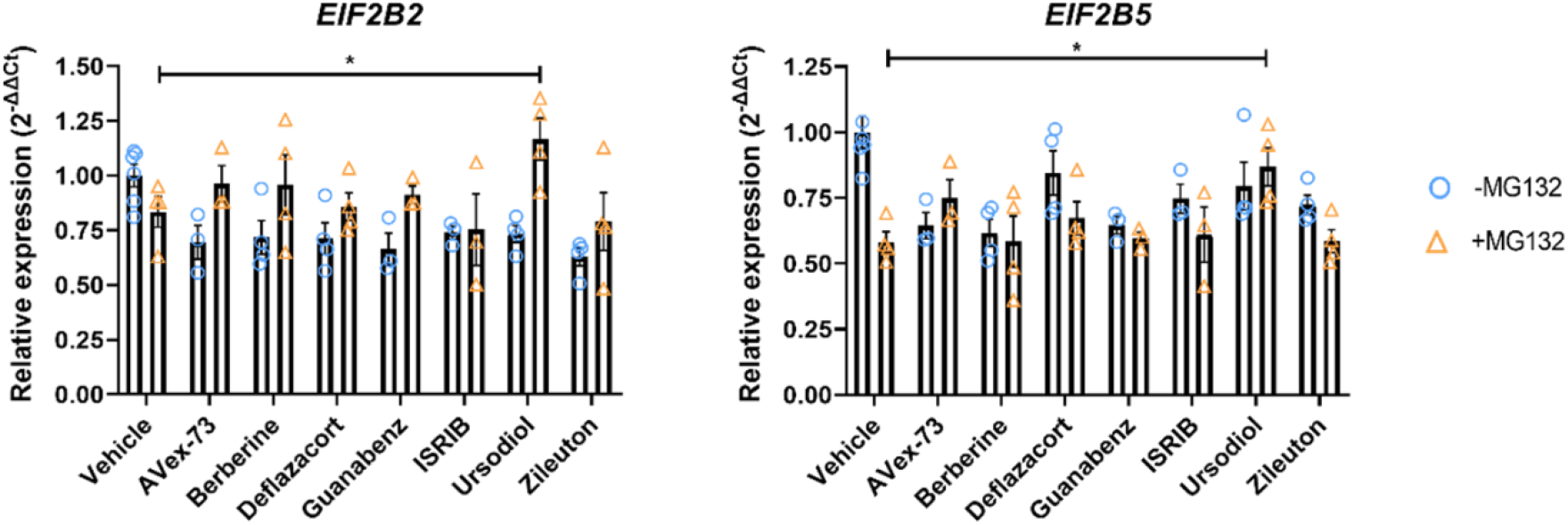
Gene expression changes of *EIF2B2* and *EIF2B5* genes in VWMD1 *EIF2B5*^R133H/A403V^ patient astrocytes treated with vehicle control (DMSO; blue circle) or 0.1 μM MG132 (orange triangle) and candidate drugs for 24 h. Relative expression was evaluated by RT-qPCR for: (A) *EIF2B2* and (B) *EIF2B5*. Individual data points are shown with mean ± SEM, n=4. Significant differences were identified by two-way ANOVA followed by Holm-Sidak posthoc test, * p < 0.05.

Together the data identified that one of the most promising drugs for clinical translation is ursodiol, a bile acid naturally formed in the liver and administered for gallstones. Ursodiol has demonstrated capacity to cross the blood brain barrier, based on clinical trials for motor neurone disease, reaching levels in cerebral spinal fluid that, based on the findings here, could be protective (38). Additionally, ursodiol has been identified as protective in other neurodegenerative and optical atrophy research (39, 40). Anti-apoptotic and neuroprotective effects of ursodiol were observed, though the underlying molecular mechanism of ursodiol-mediated protection was not identified in these studies (39, 40). Progression of VWMD includes glial cell death, developing towards neuronal cell death, and leading to paralysis and neuropathy (41). We observed ursodiol to decrease oxidative stress and possibly mitochondrial complex I burden, increase mitochondrial membrane potential, ameliorate stress-induced decreases in *EIF2B* gene expression, and improve cell viability in VWMD astrocytes under proteasomal stress. The increase of mitochondrial membrane potential by ursodiol identified in our study has also been previously observed in Alzheimer’s disease patient fibroblasts (42). Considering disrupted mitochondrial function has been extensively implicated in VWMD and a wide range of neurodegenerative disorders, further investigation of the neuroprotective effects of ursodiol are warranted. The hypothetical premise of cytoprotective drugs are those that can deliver increased cell viability, limiting disease-related degeneration. Together with information from previous studies on other CNS diseases (38, 42) our preliminary findings are supportive toward further investigation of ursodiol and anti-inflammatory drugs as cytoprotective agents for VWMD.

## Methods

### Cell culture

All experimental protocols were approved by the University of Wollongong Human Research Ethics Committee (HE17/522). Primary human dermal fibroblasts were collected from VWMD patients (2) or control family members (2) and maintained in DMEM/F12 (Life Technologies, 21331020), 10% FBS (Bovogen, SFBS-AU), 2 mM L-glutamine (Life Technologies, 25030081) and 1% penicillin/streptomycin (Life Technologies, 15140122). The mRNA-based reprogramming of patient fibroblasts into iPSCs, pluripotency marker immunofluorescence and RT-qPCR, karyotyping and mutation genotype characterisation are described in Supplementary Information (Figure S1). The iPSCs were maintained in mTESR1 (Stem Cell Technologies, 85850). Astrocytes were maintained in Astrocyte Growth Supplement Medium (AGS; Sciencell, 1852). All cells were maintained in humidified incubators at 37 °C supplemented with 5% CO_2_ for fibroblasts or hypoxic 3% O_2_ conditions for iPSCs. Neural inductions were carried out as previously described (43). Astrocyte differentiations were performed in ciliary neurotrophic factor (CNTF), epidermal growth factor (EGF) and basic fibroblast growth factor (FGF2)-based medium and transitioned to AGS, prior to characterisation by immunofluorescence and inflammatory activation, to confirm the production of functional astrocytes from iPSCs. Details of neural inductions and astrocyte differentiations are in Supplementary Information (Figure S2).

### Cell viability assay

Fibroblasts or astrocytes (5000 cells/well) were seeded in 96 well plates and incubated with a range of concentrations of hydrogen peroxide (H_2_O_2_; 0-2000 μM), MG132 (0-20 μM; Focus Bioscience, HY-13259) or thapsigargin (0-20 μM) overnight before incubation with 15 μM resazurin for 1 h and acquisition by fluorescence plate spectroscopy (excitation 544 / emission 590). For coincubation assays, cells were similarly prepared and incubated with candidate drugs and 2.5 μM MG132 (fibroblasts;) or 0.1 μM MG132 (astrocytes) for 48 h based on the IC_60_ for each cell type (Figure S3). Following incubation, cell viability was assessed by a resazurin reduction assay (Life Technologies, A13262). The coincubation drug screen was carried out in VWMD1 patient fibroblasts with the MicroSource Spectrum FDA collection (Compounds Australia), a library of 2400 FDA-approved drugs. Cell seeding and drug reagent preparation was performed by a robotic liquid handler (Hamilton Microlab Star). Single drug concentrations were selected based on protective efficacy in dose response coincubation assays and used in downstream assays at concentrations of: Annavex 2-73 (AVex-73, 5 μM), berberine (1.25 μM), guanabenz (5 μM), ISRIB (1.25 μM), deflazacort (5 μM), ursodiol (40 μM) and zileuton (1.25 μM). All treatments were performed in technical duplicate wells.

### Integrated stress response protein quantification

Fibroblasts or astrocytes were seeded in 6 well plates (200,000 cells/well) and incubated with MG132 for 0, 24 or 48 h, or coincubated with candidate compound and MG132 overnight. Cultures were lysed in RIPA buffer (50 mM Tris, pH 8.0, 150 mM NaCl, 1% Triton X-100, 0.5% Sodium Deoxycholate, 0.1% SDS and protease and phosphatase inhibitors) and stored at −20°C before sonication and quantification by a Pierce BCA Protein Assay Kit (Thermofisher Scientific, 23225). Protein lysates were heated for 5 min at 95°C in loading buffer (1% SDS, 5% 2-mercaptoethanol, 6.25% glycerol, 0.001% bromophenol blue, 0.030 M Tris-HCl, pH 6.8). Denatured protein lysates (5 μg) were loaded in 4–20% Criterion TGX Precast Gels (Bio-Rad, 5678095) and transferred to Immobilon-FL PVDF membrane (0.45 μm pore; Merck, IPFL00010). Membranes were incubated with 0.5% casein (Bio-rad, 1610782) for 1 h, primary antibody for 16 h (4 °C), and secondary antibody at 1 h (4 °C) facilitated by an automated western blot liquid handler (Cytoskeleton GOBlot). Antibodies utilised included anti-eIF2α (abcam, ab5369), anti-eIF2α-phopsho-S51 (abcam, ab32157), anti-ATF-4 (abcam, ab23760), anti-DDIT3 (abcam, ab179823), anti-GADD34 (abcam, ab126075), donkey anti-mouse Alexa Fluor Plus 488 (Thermofisher Scientific A32766) and donkey anti-rabbit Alexa Fluor Plus 647 (Thermofisher Scientific A32795). All primary antibody dilutions were performed at 1:1000, secondaries at 1:5000). Blots were imaged with a Bio-Rad ChemiDoc MP. Band intensities were normalised to stain-free total protein. Relative eIF2α phosphorylation was normalised to eIF2α. Dynamic range of signal was confirmed by linear signal to loading ratio (Figure S6).

### RT-qPCR cell stress assay

Total RNA was harvested using the PureLink RNA Mini Kit (Life Technologies, 12183025) as per manufacturer’s instructions. RNA quality was assessed using the NanoDrop 2000 Spectrophotometer (Thermofisher Scientific) with A_260/280_ ratio from 1.9 to 2.1. Up to 1 μg of RNA was used to synthesise cDNA using the iScript gDNA Clear cDNA Synthesis Kit (Bio-Rad, 1725035) and PowerUp SYBR Green (Thermofisher Scientific, A25778) was used for quantitative PCR. Primers were used at 400 nM (**Table 1).** Primer annealing temperatures were optimised by a serial dilution standard curve and considered acceptable within a range of 85-110% efficiency. RT-qPCR was performed using the QuantStudio 5 Real-Time PCR System (Applied Biosystems). Each reaction was run in triplicate and contained 20 ng of cDNA template in a final reaction volume of 20 μL. Cycling parameters were: 50 °C for 2 min, 95 °C for 2 min, then 40 cycles of 95 °C for 1 s and the annealing/extending temperature for each primer (**Table 1)** for 30 s, followed by conditions for melt curve analysis: 95 °C for 15 s, 60 °C for 1 min and 95 °C for 15 s. ΔCt values were obtained by normalisation to the average of three housekeeper genes, *GAPDH, HPRT* and *PPIA*. Data are presented using the 2^−ΔΔCt^ calculation to yield relative gene expression values (fold change).

**Table 1.**
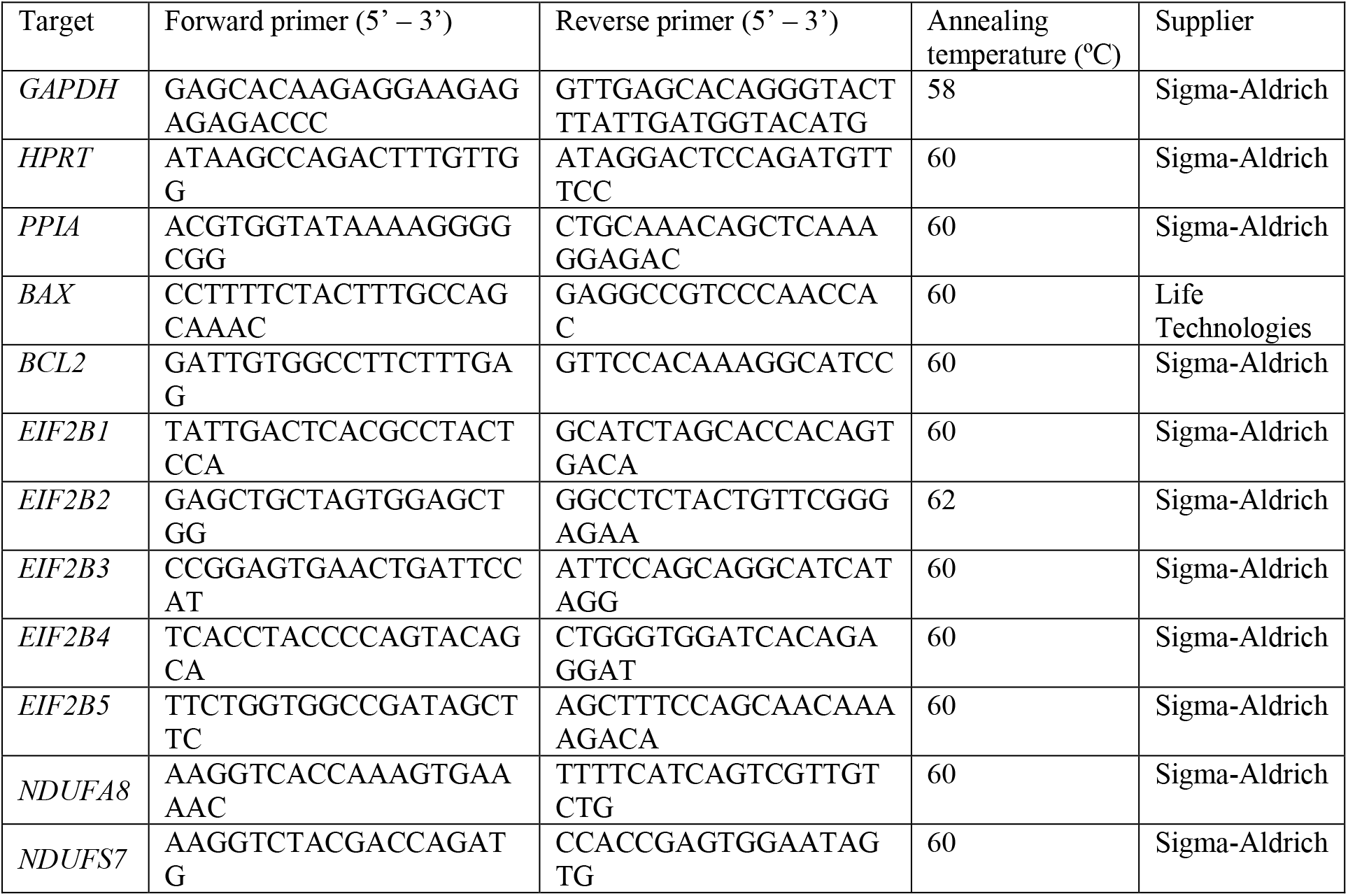
List of primers used for gene expression assay.

### Oxidative stress and mitochondrial membrane potential microscopy assays

Astrocytes (10,000 cells per well) were seeded in 96 well plates and coincubated overnight with MG132, as described above, and candidate drugs, or the positive controls for oxidative stress: H_2_O_2_ (100 μM), or the proton uncoupler carbonyl cyanide-4-phenylhydrazone (FCCP; 10 μM; Focus Bioscience, HY-100410-10MG). Cells were loaded with 20 μM H2DCFDA (DCF, Life Technologies, D399) or 0.5 μM tetramethylrhodamine ethyl ester (TMRE, Life Technologies, T669) and 10 μM Hoescht 33342 (Sigma-Aldrich, B2261-25MG) for 0.5 h before replacement of medium with phenol-red free medium and further incubation for 1 h. Relative fold change in DCF signal was acquired by fluorescence spectroscopy and normalised to cell density quantified by 0.04% sulforhodamine B (Sigma-Aldrich, 230162) solublised in 10 mM Trizma (Sigma-Aldrich, T1503) after fixation in 4% trichloroacetic acid (Sigma-Aldrich, T9159) (44). TMRE images were acquired by confocal microscopy and analysed with ImageJ to determine fluorescence normalised to cell density.

### Statistical analyses

Data are presented as the mean and standard error of the mean of at least 3 independent experiments, statistical significance was assessed by two-way analysis of variance followed by Holm-Sidak posthoc test for multiple comparisons unless otherwise stated. Significance was accepted where *p* < 0.05. Charts and statistical analyses were prepared in Prism GraphPad 8.0.

## Acknowledgements

This research was funded by philanthropic donations and a grant from the Illawarra Health and Medical Research Institute (IHMRI). L.O. is supported by a National Health and Medical Research Council (NHMRC) of Australia Boosting Dementia Research Leadership Fellowship (APP1135720). The authors thank Dr Gil Stynes (IHMRI, and Department of Surgery, Wollongong Hospital) for fibroblast collection and Prof Justin Yerbury, A/Prof Mirella Dottori and Prof Heath Ecroyd for valuable discussions, and Dr Reece Gately and Rachelle Balez for technical assistance. The authors wish to thank all of the donors, the patients and their families that have made this research possible.

## Author contributions

NN – data generation, data analysis, manuscript writing; MC – data generation, data analysis; TB – data generation, data analysis; SM – data generation, data analysis; ME - data generation; DS - data generation; DDH – data analysis; JL – data generation; SSM – data analysis; NSB – data generation; CS – data generation; LO - intellectual input, data analysis, manuscript writing, supervision, funding. All authors reviewed the manuscript.

## Conflict of Interest

The authors declare no conflict of interest.

## Notes

### Competing Interest Statement

The authors have declared no competing interest.

## References

1. Bugiani M, Vuong C, Breur M, van der Knaap MS. Vanishing white matter: a leukodystrophy due to astrocytic dysfunction. 2018;28(3):408–21.

2. Moon SL, Parker R. EIF2B2 mutations in vanishing white matter disease hypersuppress translation and delay recovery during the integrated stress response. RNA. 2018;24(6):841–52.

3. Abbink TEM, Wisse LE, Jaku E, Thiecke MJ, Voltolini-González D, Fritsen H, et al. Vanishing white matter: deregulated integrated stress response as therapy target. Annals of clinical and translational neurology. 2019;6(8):1407–22.

4. Sidrauski C, Tsai JC, Kampmann M, Hearn BR, Vedantham P, Jaishankar P, et al. Pharmacological dimerization and activation of the exchange factor eIF2B antagonizes the integrated stress response. elife. 2015;4:e07314.

5. Brush MH, Shenolikar S. Control of cellular GADD34 levels by the 26S proteasome. Molecular cellular biology. 2008;28(23):6989–7000.

6. Dooves S, Bugiani M, Postma NL, Polder E, Land N, Horan ST, et al. Astrocytes are central in the pathomechanisms of vanishing white matter. The Journal of Clinical Investigation. 2016;126(4):1512–24.

7. Leferink PS, Dooves S, Hillen AEJ, Watanabe K, Jacobs G, Gasparotto L, et al. Astrocyte Subtype Vulnerability in Stem Cell Models of Vanishing White Matter. Annals of Neurology. 2019;86(5):780–92.

8. Zhou L, Li P, Chen N, Dai L-F, Gao K, Liu Y-N, et al. Modeling vanishing white matter disease with patient-derived induced pluripotent stem cells reveals astrocytic dysfunction. CNS Neuroscience & Therapeutics. 2019;25(6):759–71.

9. Atzmon A, Herrero M, Sharet-Eshed R, Gilad Y, Senderowitz H, Elroy-Stein O. Drug screening identifies sigma-1-receptor as a target for the therapy of VWM leukodystrophy. Frontiers in molecular neuroscience. 2018;11:336.

10. Wong YL, LeBon L, Basso AM, Kohlhaas KL, Nikkel AL, Robb HM, et al. eIF2B activator prevents neurological defects caused by a chronic integrated stress response. ELife. 2019;8:e42940.

11. Weng T-Y, Tsai S-YA, Su T-P. Roles of sigma-1 receptors on mitochondrial functions relevant to neurodegenerative diseases. Journal of biomedical science. 2017;24(1):74.

12. Park WH, Kim SH. MG132, a proteasome inhibitor, induces human pulmonary fibroblast cell death via increasing ROS levels and GSH depletion. J Oncology reports. 2012;27(4):1284–91.

13. Jiang H-Y, Wek RC. Phosphorylation of the α-subunit of the eukaryotic initiation factor-2 (eIF2α) reduces protein synthesis and enhances apoptosis in response to proteasome inhibition. Journal of Biological Chemistry. 2005;280(14):14189–202.

14. Brush MH, Weiser DC, Shenolikar S. Growth arrest and DNA damage-inducible protein GADD34 targets protein phosphatase 1 alpha to the endoplasmic reticulum and promotes dephosphorylation of the alpha subunit of eukaryotic translation initiation factor 2. Molecular and cellular biology. 2003;23(4):1292–303.

15. Teske BF, Fusakio ME, Zhou D, Shan J, McClintick JN, Kilberg MS, et al. CHOP induces activating transcription factor 5 (ATF5) to trigger apoptosis in response to perturbations in protein homeostasis. Mol Biol Cell. 2013;24(15):2477–90.

16. Garcia-Segura LM. Hormones and brain plasticity: Oxford University Press; 2009.

17. Turón-Viñas E, Pineda M, Cusí V, López-Laso E, Del Pozo RL, Gutiérrez-Solana LG, et al. Vanishing white matter disease in a spanish population. Journal of central nervous system disease. 2014;6:JCNSD. S13540.

18. Petnikota H, Madhuri V, Gangadharan S, Agarwal I, Antonisamy B. Retrospective cohort study comparing the efficacy of prednisolone and deflazacort in children with muscular dystrophy: A 6 years’ experience in a South Indian teaching hospital. Indian J Orthop. 2016;50(5):551–7.

19. McAdam LC, Mayo AL, Alman BA, Biggar WD. The Canadian experience with long-term deflazacort treatment in Duchenne muscular dystrophy. Acta Myol. 2012;31(1):16–20.

20. Kawasaki T, Kitao T, Nakagawa K, Fujisaki H, Takegawa Y, Koda K, et al. Nitric oxide-induced apoptosis in cultured rat astrocytes: Protection by edaravone, a radical scavenger. Glia. 2007;55(13):1325–33.

21. Sato T, Mizuno K, Ishii F. A novel administration route of edaravone–II: mucosal absorption of edaravone from edaravone/hydroxypropyl-beta-cyclodextrin complex solution including L-cysteine and sodium hydrogen sulfite. Pharmacology. 2010;85(2):88–94.

22. Parikh A, Kathawala K, Tan CC, Garg S, Zhou X-F. Development of a novel oral delivery system of edaravone for enhancing bioavailability. International journal of pharmaceutics. 2016;515(1-2):490–500.

23. Rothstein JD. Edaravone: a new drug approved for ALS. Cell. 2017;171(4):725.

24. Wu Y, Pan Y, Du L, Wang J, Gu Q, Gao Z, et al. Identification of novel EIF2B mutations in Chinese patients with vanishing white matter disease. Journal of human genetics. 2009;54(2):74–7.

25. Ortori C, Barrett D, Fischer P, Halliday M, Radford H, Sekine Y, et al. Partial restoration of protein synthesis rates by the small molecule ISRIB prevents neurodegeneration without pancreatic toxicity. 2015.

26. Sidrauski C, Acosta-Alvear D, Khoutorsky A, Vedantham P, Hearn BR, Li H, et al. Pharmacological brake-release of mRNA translation enhances cognitive memory. Elife. 2013;2:e00498.

27. Palam L, Gore J, Craven K, Wilson J, Korc M. Integrated stress response is critical for gemcitabine resistance in pancreatic ductal adenocarcinoma. Cell death & disease. 2015;6(10):e1913–e.

28. Nguyen HG, Conn CS, Kye Y, Xue L, Forester CM, Cowan JE, et al. Development of a stress response therapy targeting aggressive prostate cancer. Science translational medicine. 2018;10(439):eaar2036.

29. Wang L, Popko B, Tixier E, Roos RP. Guanabenz, which enhances the unfolded protein response, ameliorates mutant SOD1-induced amyotrophic lateral sclerosis. Neurobiology of disease. 2014;71:317–24.

30. Halliday M, Radford H, Sekine Y, Moreno J, Verity N, Le Quesne J, et al. Partial restoration of protein synthesis rates by the small molecule ISRIB prevents neurodegeneration without pancreatic toxicity. Cell death & disease. 2015;6(3):e1672–e.

31. Wong YL, LeBon L, Edalji R, Lim HB, Sun C, Sidrauski C. The small molecule ISRIB rescues the stability and activity of Vanishing White Matter Disease eIF2B mutant complexes. Elife. 2018;7:e32733.

32. Vuddanda PR, Chakraborty S, Singh S. Berberine: a potential phytochemical with multispectrum therapeutic activities. Expert opinion on investigational drugs. 2010;19(10):1297–307.

33. Yu W, Sheng M, Xu R, Yu J, Cui K, Tong J, et al. Berberine protects human renal proximal tubular cells from hypoxia/reoxygenation injury via inhibiting endoplasmic reticulum and mitochondrial stress pathways. Journal of translational medicine. 2013;11(1):24.

34. Li Z, Geng Y-N, Jiang J-D, Kong W-J. Antioxidant and anti-inflammatory activities of berberine in the treatment of diabetes mellitus. Evidence-Based Complementary Alternative Medicine. 2014;2014.

35. Raini G, Sharet R, Herrero M, Atzmon A, Shenoy A, Geiger T, et al. Mutant eIF2B leads to impaired mitochondrial oxidative phosphorylation in vanishing white matter disease. 2017;141(5):694–707.

36. Guiraud S, Davies KE. Pharmacological advances for treatment in Duchenne muscular dystrophy. Current opinion in pharmacology. 2017;34:36–48.

37. Liu Y, Wang W, Li Y, Xiao Y, Cheng J, Jia J. The 5-lipoxygenase inhibitor zileuton confers neuroprotection against glutamate oxidative damage by inhibiting ferroptosis. Biological Pharmaceutical Bulletin. 2015;38(8):1234–9.

38. Parry GJ, Rodrigues CM, Aranha MM, Hilbert SJ, Davey C, Kelkar P, et al. Safety, tolerability, and cerebrospinal fluid penetration of ursodeoxycholic acid in patients with amyotrophic lateral sclerosis. Journal of Clinical Neuropharmacology. 2010;33(1):17–21.

39. Boatright JH, Nickerson JM, Moring AG, Pardue MT. Bile acids in treatment of ocular disease. Journal of ocular biology. 2009;2(3):149.

40. Foster SL, Kendall C, Lindsay AK, Ziesel AC, Allen RS, Mosley SS, et al. Development of Bile Acids as Anti-Apoptotic and Neuroprotective Agents in Treatment of Ocular Disease. Drug Product Development for the Back of the Eye: Springer; 2011. p. 565–76.

41. Barros SR, Parreira SC, Miranda AF, Pereira AM, Campos NM. New insights in vanishing white matter disease: Isolated bilateral optic neuropathy in adult onset disease. Journal of Neuro-Ophthalmology. 2018;38(1):42–6.

42. Bell SM, Barnes K, Clemmens H, Al-Rafiah AR, Al-ofi EA, Leech V, et al. Ursodeoxycholic acid improves mitochondrial function and redistributes Drp1 in fibroblasts from patients with either sporadic or familial Alzheimer’s disease. Journal of molecular biology. 2018;430(21):3942–53.

43. Denham M, Dottori M. Neural differentiation of induced pluripotent stem cells. Neurodegeneration: Springer; 2011. p. 99–110.

44. Vichai V, Kirtikara K. Sulforhodamine B colorimetric assay for cytotoxicity screening. Nature protocols. 2006;1(3):1112.

