## Supplementary data for "Identification of repurposable cytoprotective drugs for Vanishing White Matter Disease"

### Supplementary Information

#### iPSC generation

Dermal fibroblasts were isolated from skin biopsies of VWMD patients and non-disease individuals approved by the University of Wollongong Human Ethics Committee. Control lines were gender matched; the VWMD1 control line was generated from a healthy carrier family member of VWMD1 and the VWMD6 control line was created from a family member of VWMD6 bearing one G200V mutant allele.  Fibroblasts were reprogrammed with by a non-integrating method following the manufacturer's instructions (Reprocell, 00-0076). Fibroblasts from passage 1-2 were seeded at several densities on laminin 511 (iMatrix, Reprocell, NP892-011) coated plates and transfected with the RNA plus micro-RNA mix using the Lipofectamine RNAiMAX Transfection Reagent (Thermofisher Scientific, 13778075) for four days consecutively. Fresh PSC maintenance medium (mTeSR-1, StemCell Technologies, 85850) was changed daily. Primary iPSC colonies were manually selected after day 10. Established iPSC lines were maintained in mTeSR-1 on matrigel-coated dishes at 37 °C and 5% CO2 and 3% O2 and passaged in PBS/EDTA.

**Sequencing.** Genomic DNA was isolated from fibroblasts or iPSCs with the PureLink RNA Mini Kit (Life Technologies, 12183018A) or Isolate II Genomic DNA kit (Bioline, BIO-52075) and the relevant regions of interest PCR-amplified using PrimeStar HS DNA polymerase (Takara, R010B). *EIF2B5* genotyping primers were designed within introns generate PCR products spanning c.338G>A, p.Arg113His (345 bp) and c.1208C>T, p.Ala403Val (289 bp). *EIF2B2* genotyping primers were designed within introns to generate PCR products spanning c.599G>T, p. Gly200Val and c.638A>G, p.Glu213Gly (292 bp) (Table S1).

**Karyotyping.** Karyotyping was performed at Sullivan Nicolaides Pathology Pty Ltd (Brisbane, Australia), on iPSCs between passages 8 and 11, with 15 metaphase spreads counted at 400 bphs.

**Immunofluorescence.** Samples from each iPSC line were seeded on glass coverslips and fixed with 4% paraformaldehyde for 10 minutes. Cells were permeabilised with a 0.05% Triton-X (Sigma-Aldrich, T8787) solution and blocked with 10% goat serum (Life Technologies, 16210-064) before overnight incubation with anti-Oct4 (StemCell Technologies, 60059, 1:1000) or anti-SSEA4 (Abcam, ab16287, 1:200) antibodies, followed by 1 h incubation with secondary antibody (Life Technologies, A11008) and Hoechst 33342 (Life Technologies, H3570).

**qPCR.** Fibroblasts and iPSCs RNA was extracted column-free using the TRIsure reagent (Bioline, BIO-38033) followed by cDNA synthesis (Tetro cDNA Synthesis Kit, Bioline, BIO-65043), both as per manufacture's protocol. Expression of *POU5F1* and *NANOG* were assessed by qPCR using the SensiFAST SYBR No-ROX Kit master mix (Bioline, BIO-98020) (Table S1). H9 and the source fibroblast for reference and the H9 expression levels were set to 1 for comparison. *GAPDH* and *HPRT1* were used as housekeeping for normalisation of the data. Relative expression was calculated by the ΔΔCt method.

**Scorecard.** iPSCs were directed to each germ layer separately. Mesoderm and endoderm differentiation was achieved by using STEMdiff Mesoderm Induction Medium (StemCell Technologies, 05221) and STEMdiff Definitive Endoderm Kit (StemCell Technologies, 05110), respectively, as per manufacture's protocol. Differentiation towards the ectodermal germ layer was carried out in 50% DMEM/F12 (Gibco, 12800082), 50% Neurobasal (Gibco, 21103-049) supplemented with N2 (1x, Gibco, 17502001), B27 (1x, Gibco, 12587-010), 1x ITS-A (Gibco, 51300-044), L-Glutamine (2 mM, Gibco, 25030-081) and D-(+)-Glucose (0.3%, Sigma Aldrich, G8769) in the presence of SB-431542 (10 µM, MedChemExpress, HY-10431), CHIR-99021 (3µM, MedChemExpress, HY-10182B) and LDN193189 (100 nM, MedChemExpress, HY-12071A) for five days. Equal amounts of cDNA from each germ layer (1:1:1) were pooled together to make up a total of one microgram of cDNA, required for the ScoreCard (TaqMan hPSC Scorecard, A15876). The ScoreCard was run according to manufacture's instructions.

**Table S1. Primers used for genotyping and pluripotency markers.**

| Target | Forward primer | Reverse primer | Annealing temperature (ºC) | Supplier |
|---|---|---|---|---|
| *EIF2B5 R113H* | CCATCGAGAAGGACTGTG | GTCTCAGGGCTCTGCTG | 56 | Sigma-Aldrich |
| *EIF2B5 A403V* | GGTCTTCCCATCCTGAGC | GTGCATCCTGATAATGAGAGC | 54 | Sigma-Aldrich |
| *EIF2B2 G200V & E213G* | GTGCTGGATATGCCCATATC | CACCGTGGATATTACAGGAG | 55 | Sigma-Aldrich |
| *POU5F1* | GATCACCCTGGGATATACAC | GCTTTGCATATCTCCTGAAG | 58 | Sigma-Aldrich |
| *NANOG* | CCAGAACCAGAGAATGAAATC | TGGTGGTAGGAAGAGTAAAG | 58 | Sigma-Aldrich |
| *GAPDH* | GAGCACAAGAGGAAGAGAGAGACCC | GTTGAGCACAGGGTACTTTATTGATGGTACATG | 58 | Sigma-Aldrich |
| *HPRT1* | TGACACTGGCAAAACAATGCA | GGTCCTTTTCACCAGCAAGCT | 58 | Life Technologies |

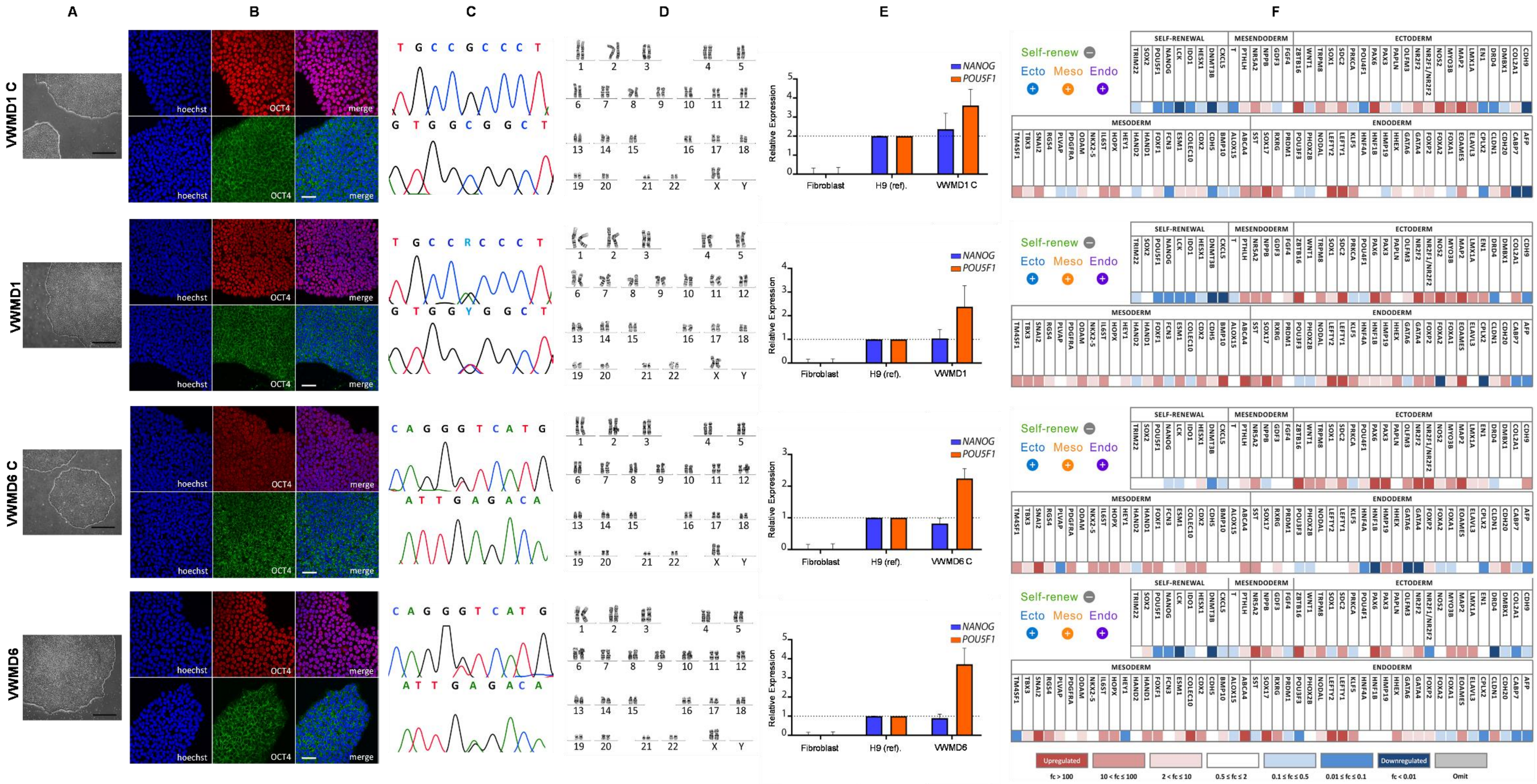
A
B
C
D
E
F
VWMD1 C
VWMD1
VWMD6 C
VWMD6
hoechst
OCT4
merge
T G C C G C C C T
G T G G C G G C T
T G C C R C C C T
G T G G Y G G C T
C A G G G T C A T G
A T T G A G A C A
1 2 3 4 5
6 7 8 9 10 11 12
13 14 15 16 17 18
19 20 21 22 X Y
NANOG
POU5F1
Relative Expression
Fibroblast
H9 (ref).
VWMD1 C
VWMD1
VWMD6 C
VWMD6
Self-renew
Ecto Meso Endo
SELF-RENEWAL
MESENDODERM
ECTODERM
MESODERM
ENDODERM
Upregulated
Downregulated
fc > 100
10 < fc ≤ 100
2 < fc ≤ 10
0.5 ≤ fc ≤ 2
0.1 ≤ fc ≤ 0.5
0.01 ≤ fc ≤ 0.1
fc < 0.01
Omit

**Figure S1. iPSC characterisation for: VWMD1 control (VWMD1 C; iPSC line clone 1) from healthy non-carrier control, gender-matched, donor related to VWMD1; VWMD1 (iPSC line clone 3) from *EIF2B5*$^{R113H/A403V}$ patient; VWMD6 control (VWMD6 C; iPSC line clone 2) from healthy non-disease carrier control (*EIF2B2*$^{G200V}$), gender-matched, donor related to VWMD6; VWMD6 (iPSC line clone 2) from *EIF2B2*$^{G200V/E213G}$ patient. (A) Phase contrast images of representative iPSC colonies (scale bar = 200 µm). (B) Immunofluorescence images of OCT4 (red) and SSEA (green) pluripotency markers, with Hoechst 33342 nuclear stain (blue) (scale bar = 50 µm). (C) Chromatogram of VWMD1 Control *EIF2B5*$^{wt/wt}$ sequence encoding wild-type R113 (top panel) and A403 (bottom panel); for VWMD1 patient *EIF2B5*$^{R113H/A403V}$ sequence encoding R113H (top panel) and A403V (bottom panel) mutations; VWMD6 Control *EIF2B2*$^{G200V/wt}$ non-disease control carrier sequence encoding one mutant allele G200V (top panel) and one wild-type allele E213 (bottom panel); VWMD6 from *EIF2B2*$^{G200V/E213G}$ patient sequence encoding G200V (top panel) and E213G (bottom panel). (D) Karyotype analysis confirming normal karyotype of reprogrammed cells. (E) Quantitative reverse transcription PCR confirming upregulation of *NANOG* and *POU5F1* in iPSCs, at similar levels to the pluripotent embryonic stem cell line H9. (F). Scorecard characterisation confirms reprogrammed cells are pluripotent and able to differentiate into cells from the three germ layers, endoderm, mesoderm and ectoderm. Shown are heat maps representing the fold upregulation of differentiated genes and downregulation of genes involved in self-renewal (stem cell markers) upon differentiation.**

**Neural induction and astrocyte differentiations**

iPSC colonies were incubated with neural induction media, which consisted of 2% B-27 (Life Technologies, 17504044), 1% N-2 (Life Technologies, 17502001), 2 mM L-glutamine, 10 µM SB431542 (Focus Bioscience, HY-10431) and 100 nM LDN193189 (Focus Bioscience, HY-12071A) for 7 days, before replacement of small molecules with 20 ng/mL EGF (StemCell Technologies, 78006) and 20 ng/mL FGF-2 (Miltenyi Biotec, 130-093-564) for an additional 7 days. Neural rosettes were triturated and collected, and NPCs were expanded with subsequent passages and cryopreserved. For astrocyte differentiations, NPCs were seeded in 175 cm2 flasks (20000 cells/$cm^2$) and routinely subcultured in DMEM/F12 supplemented with 1% B27, 0.1% N2, 5 ng/mL CNTF (Miltenyi Biotec, 130-108-972), 5 ng/mL EGF, 2 ng/mL FGF2, 2% FBS, 0.2 mM ascorbic acid, 2 mM L-glutamine for 21 days, before replacement of medium with DMEM/F12 supplemented with Astrocyte Growth Supplement (Sciencell, 1852).

**Neural cell immunofluorescence characterisation**

Neural stem cells (20,000 cells per well) or astrocytes (10,000 cells per well) were seeded in duplicate wells overnight in microscopy grade 96 well microtitre plates and fixed for 10 min in 4% paraformaldehyde. Samples were incubated with blocking buffer (5% donkey serum, 0.05% Triton X-100, 0.05% Tween-20) incubated with primary antibodies overnight at $4^oC$, washed 3 times in (PBS 0.05% Tween 20) for 5 min, incubated with secondary antibody for 1 h, washed, and incubated with SYTOX Blue and 0.1 mg/mL RNAse. Antibodies used in this study were anti-SOX2 (Thermofisher, PA1-094), anti-nestin (Thermofisher, MA1-110, 1:300), anti-ALDOC (abcam, ab190368, 1:300), anti-Aquaporin 4 (abcam, ab9512, 1:300), anti-EAAT1 (abcam, ab416, 1:300), anti-EAAT2 (abcam, ab41621, 1:300), anti-GFAP (abcam, ab53554, 1:300), donkey anti-mouse AlexaFluor 488 (abcam, ab150105, 1:500), donkey anti-goat AlexaFluor 568 (abcam, ab175704, 1:500), and donkey anti-rabbit AlexaFluor 647 (abcam, ab150075, 1:500). All samples were counterstained with 10 µM Hoescht 33342 for 10 min. Antibody validation was performed by staining on rat hippocampal dentate gyrus regions (Figure S6). >300 cells were quantified by confocal microscopy (Leica SP8) with a 20X objective.

**Astrocyte inflammation assay**

Cells were seeded in 12 well plates (25000 cells/cm$^2$) before overnight incubation with polyinosinic:polycytidylic acid (10 µg/mL) in duplicate wells. Supernatants were collected, centrifuged at 3000 × g, to remove debris prior to storage at -20 $^o$C until analysed. Supernatants were analysed by ELISA kits for RANTES (Thermofisher Scientific EHRNTS) and IL-6 (Thermofisher Scientific EH2IL6).

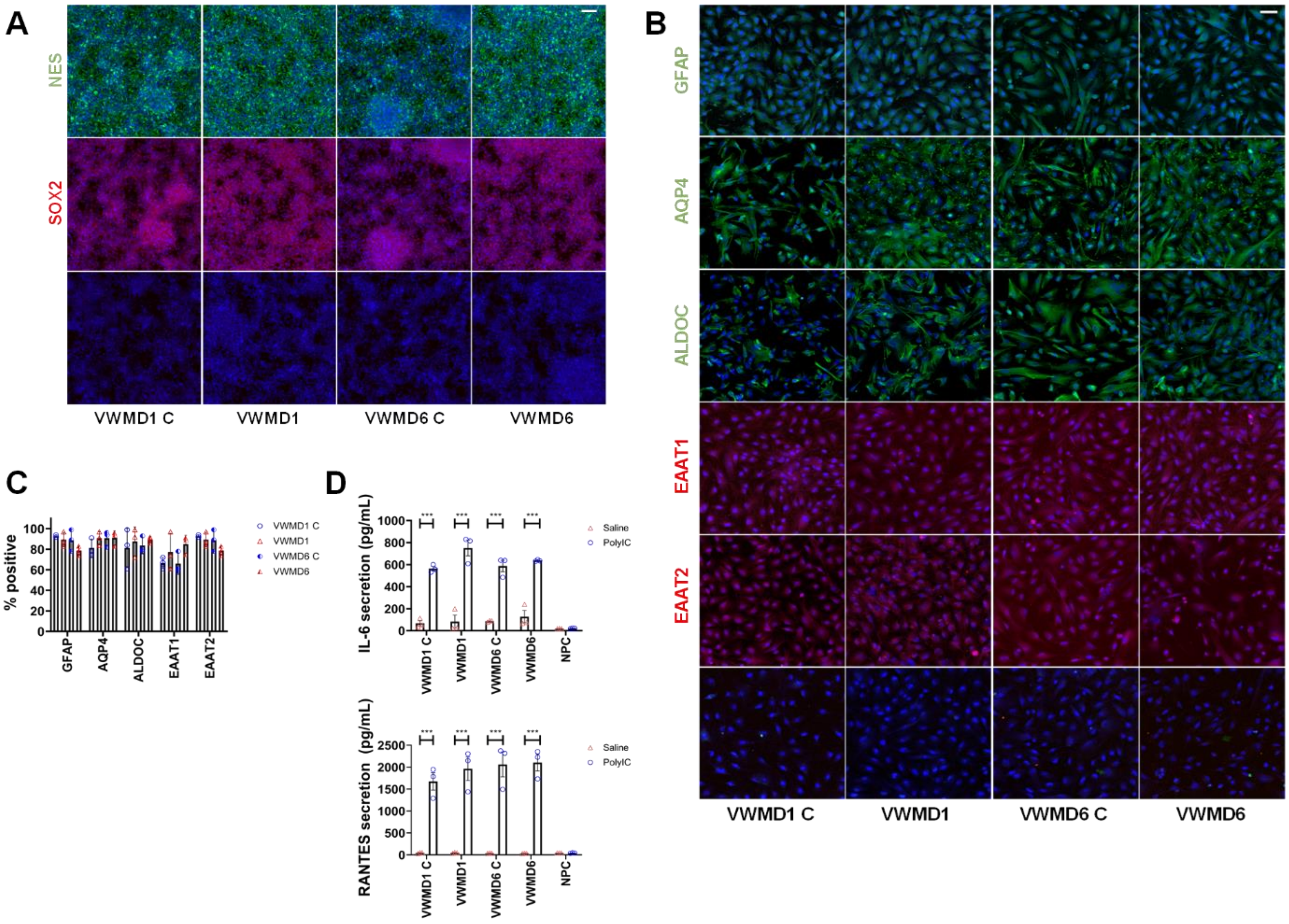


**Figure S2. Neural induction and astrocyte differentiation immunofluorescence and immunoreactivity from iPSCs. (A) Neural stem cell nestin and SOX2 immunofluorescence (n = 3); bottom row no primary control. (B) Representative images of astrocyte markers GFAP, AQP4, ALDOC, EAAT1, EAAT2 immunofluorescence (scale bar = 30 µm, bottom row no primary control). (C) Quantification of immunofluorescence astrocyte marker expression in individual lines (%), n=3 independent replicates; individual data points and mean ± SEM shown. (D) IL-6 and RANTES secretion in iPSC-derived astrocytes in presence of saline (vehicle control) or polyinosinic:polycytidylic acid (polyIC) inflammatory stimulation measured by ELISA, neural precursor cells (NPC) used as a negative control; n = 3; individual data points and mean ± SEM shown; significant differences identified by two-way ANOVA with Holm-Sidak posthoc test, *** $p < 0.001$.**

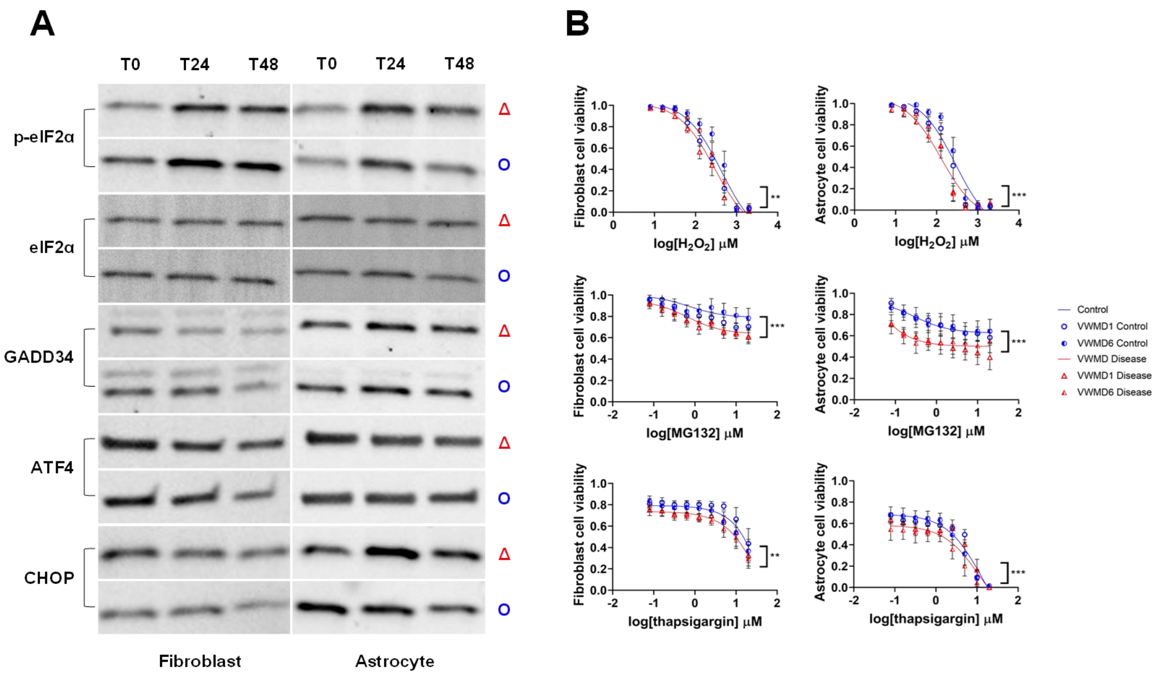


**Figure S3. VWMD ISR marker expression and cell viability under stress. (A) Representative western blots of phosphorylated eIF2α (~37 kDa), eIF2α (~37 kDa), GADD34 (~50 kDa), ATF4 (~76 kDa) and CHOP (~26 kDa) expression in VWMD1 patient fibroblasts and VWMD1 iPSC-derived astrocytes under MG132 stress at 0, 24 and 48 h. (B) Relative cell viability of VWMD patient disease and control lines treated with a range of concentrations of $H_2O_2$, MG132 and thapsigargin for 48 h. Individual data points are shown for Control cell lines (blue circles) and VWMD cell lines (red triangles) with mean ± SEM, n=7; significant differences were identified between control and VWMD by two-way ANOVA with Holm-Sidak posthoc test, *** $p < 0.001$.**

**Table S2.** Candidate drug panel and classes; primary screen score in fibroblasts.

| Drug name | Class | Primary screen score |
|---|---|---|
| Amitriptyline | Sigma-1 receptor agonist | - |
| AVex-73 | Sigma-1 receptor agonist | - |
| Berberine | Alkaloid | - |
| Budesonide | Glucocorticosteroid | 1.40 |
| Curcumin | Antioxidant | 1.22 |
| Deferasirox | Iron chelator | 1.30 |
| Deferoxamine | Iron chelator | 1.20 |
| Deflazacort | Glucocorticosteroid | 1.37 |
| Dexamethasone | Glucocorticosteroid | 1.35 |
| Edaravone | Antioxidant | 1.22 |
| Guanabenz | Integrated stress response enhancement | - |
| Hydrocortisone | Glucocorticosteroid | 1.21 |
| ISRIB | Integrated stress response inhibitor | - |
| Methylprednisolone | Glucocorticosteroid | 1.33 |
| Prednisolone | Glucocorticosteroid | 1.23 |
| Probucol | Antioxidant | 1.15 |
| Tauroursodeoxycholate | Bile acid | - |
| Triamcinolone | Glucocorticosteroid | 1.17 |
| Ursodiol | Bile acid | 1.15 |
| Zileuton | 5-lipooxygenase inhibitor | 1.32 |

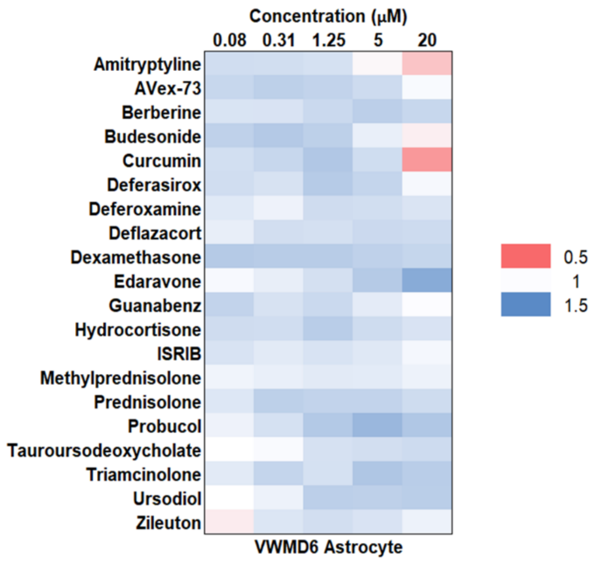


**Figure S4. VWMD6 *EIF2B2*$^{G200V/E213G}$ patient iPSC-derived astrocyte cell line candidate drug panel, ISR and EIF2B gene expression changes. (A) Heat map represents cytoprotection (blue) versus cytotoxicity (red) of individual drugs at increasing concentrations in the presence of MG132 in VWMD6 astrocytes; cell viability changes relative to MG132 stress (n=5-6).**

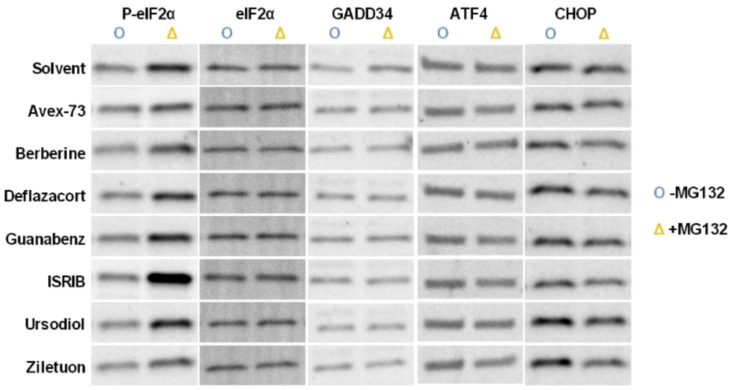


**Figure S5. Representative western blots of ISR markers in VWMD1 iPSC-derived astrocytes. Phosphorylated eIF2α (~37 kDa), eIF2α (~37 kDa), GADD34 (~50 kDa), ATF4 (~76 kDa) and CHOP (~26 kDa) expression in vehicle control (DMSO, blue circle) or with 0.1 µM MG132 (orange triangle) with candidate drugs.**

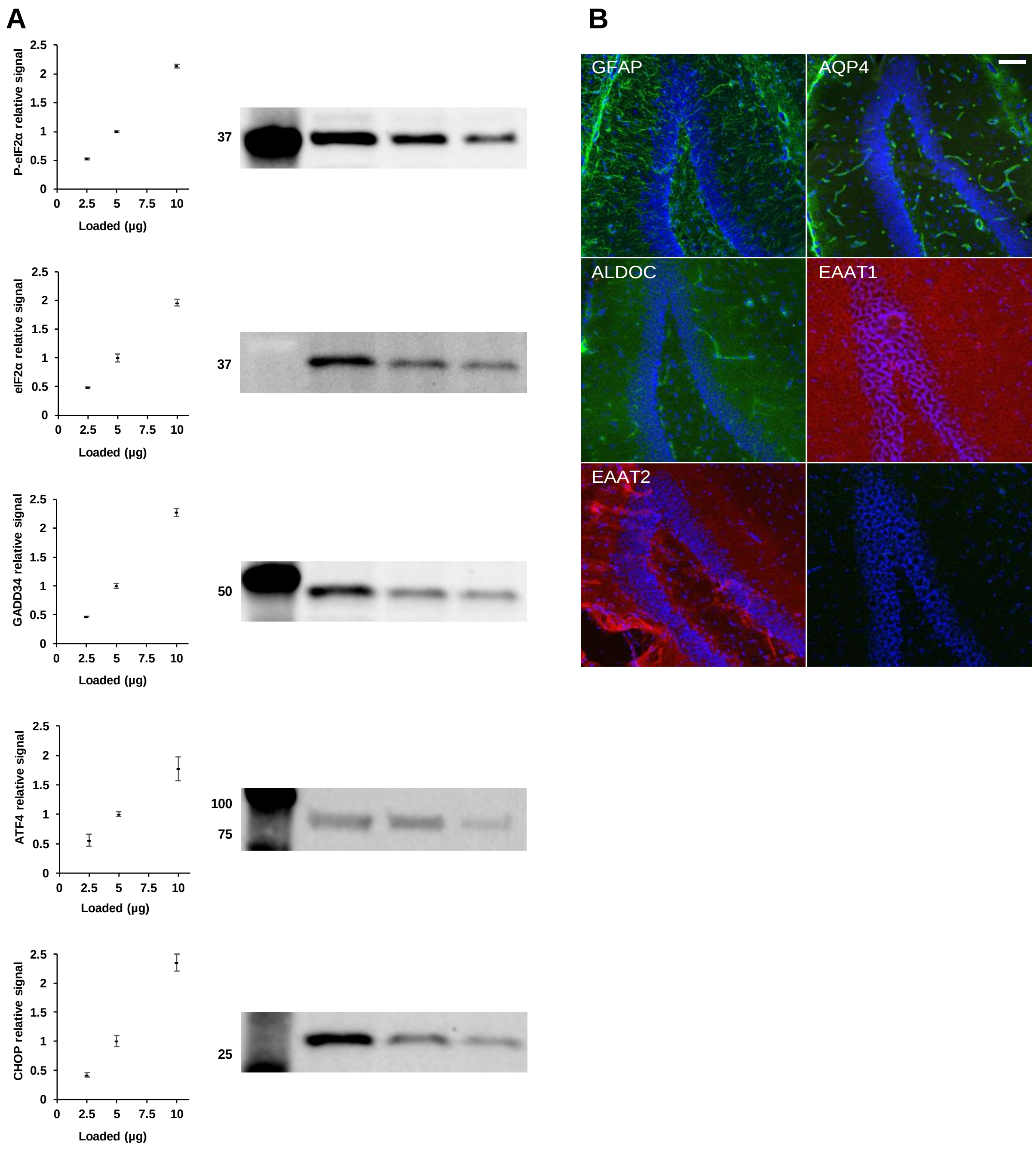


**Figure S6. Antibody validation in semi-quantitative western blots and astrocyte immunofluorescence characterisation panel. (A) Representative blots demonstrate proportional signal change to loading of ISR markers with antibodies to phosphorylated eIF2α, eIF2α, GADD34, ATF4 and CHOP in VWMD1 control astrocytes (data points are mean ± SEM, n = 3). (B) Expected regions of staining of rat hippocampal cryoslices around the dentate gyrus (scale bar = 60 μm) with antibodies to GFAP, AQP4, ALDOC, EAAT1, EAAT2 or no primary control.**
